# A Charcot–Marie–Tooth–associated dynamin mutation causes peripheral neuropathy-like phenotypes via Patronin/CAMSAP-dependent microtubule stabilization

**DOI:** 10.64898/2026.11.10.765802

**Authors:** Yuki Miki, Misako Okumura, Takahiro Chihara, Kozue Hamao

## Abstract

Dynamin-2 mutations are associated with Charcot–Marie–Tooth (CMT) neuropathy; however, how these mutations affect neuronal function *in vivo* remains unclear. Here, we generated a *Drosophila* model carrying a CRISPR/Cas9-mediated knock-in of the *shi^549Δ3^* mutation, corresponding to the human dynamin-2-551Δ3 variant, and examined its physiological and cellular consequences. The *shi^549Δ3^* mutants exhibited progressive motor impairment across multiple genetic conditions. At the neuromuscular junction, the mutation caused synaptic overgrowth accompanied by excessive microtubule stabilization, including increased Futsch-positive loop structures and bouton numbers. Genetic interaction analysis revealed that reduction of Patronin, the *Drosophila* homolog of mammalian CAMSAPs, partially suppressed lethality and motor defects, linking microtubule stabilization to neuronal dysfunction *in vivo*. Consistently, CAMSAP2 knockdown in mammalian cells attenuated dynamin-2-induced microtubule acetylation, supporting a conserved mechanism. Together, these findings identify dysregulated microtubule stabilization as a key mechanism underlying neuropathy-like phenotypes caused by a CMT-associated dynamin mutation *in vivo*.

## INTRODUCTION

Charcot–Marie–Tooth disease (CMT) is the most common inherited peripheral neuropathy, affecting approximately 1 in 2,500 individuals ^1,2^. It is characterized by progressive, length-dependent degeneration of motor and sensory nerves, with clinical symptoms including distal muscle weakness and atrophy, foot deformities, steppage gait, distal sensory loss, and mild to severe neuropathic pain ^1–3^.

CMT is classified into three groups: type 1 CMT (CMT1; characterized mainly by demyelination), type 2 CMT (CMT2; characterized mainly by axonal degeneration), and dominant intermediate forms (DI-CMT), which exhibit features of both demyelination and axonal degeneration ^4,5^. CMT2 is further subdivided into multiple subtypes, defined by the causative genes. More than 100 disease-associated genes have been identified, including the dynamin-2 gene (*DNM2*) mutations in which are linked to CMT2M and DI-CMTB ^6,7^.

Dynamin was originally identified as a microtubule-binding protein ^8–11;^ however, most functional studies have focused on its well-established role in endocytosis. Dynamin is evolutionarily conserved across diverse animals, including insects (e.g., *Drosophila melanogaster*) ^12,13^, nematodes (e.g., *Caenorhabditis elegans*) ^13^, and mammals ^14^. In *D. melanogaster*, dynamin is encoded by the *shibire* (*shi*) gene, whereas mammals express three dynamins: dynamin-1, dynamin-2, and dynamin-3, encoded by *DNM1*, *DNM2*, and *DNM3*, respectively. Dynamin-1 is enriched in the nervous system, dynamin-2 is ubiquitously expressed, and dynamin-3 is expressed in several tissues, including the brain, heart, lungs, and testes ^15,16^.

Dynamin contains five domains: the GTPase, middle, pleckstrin homology, GTPase effector, and proline-rich domains. GTPase activity is essential for membrane fission during endocytosis ^17^. In addition, dynamin interacts directly with microtubules, forming bundles *in vitro* ^9,10^ and exhibiting increased GTPase activity upon polymerization around microtubules ^18^. A deletion of three amino acids in the pleckstrin homology domain of dynamin-2 (D551_E553del), identified in patients with DI-CMTB in which it is typically present in the heterozygous state, is commonly referred to as dynamin-2-551Δ3 and corresponds to dynamin-2-555Δ3, based on the longest isoform numbering ^6^. Overexpression of dynamin-2-551Δ3 increases microtubule acetylation, a marker of microtubule stabilization, in cultured cells. Similarly, depletion of dynamin-2 induces microtubule stabilization and also decreases the dynamic instability of microtubules and impairs microtubule-dependent vesicular transport ^19^. These observations indicate that dynamin-2-551Δ3 perturbs normal microtubule regulation, although the underlying mechanism remains unclear.

Our previous work demonstrated that the self-assembly activity of dynamin-2, rather than its GTPase activity, is required for regulating microtubule stability, indicating that the mechanisms underlying microtubule regulation and endocytosis are distinct ^20^. However, the mechanisms underlying how disease-associated dynamin-2 mutations affect neuronal structure and function *in vivo*, and how excessive microtubule stabilization contributes to CMT pathogenesis, remain poorly understood.

In this study, we generated a *Drosophila* model carrying a CRISPR/Cas9-mediated knock-in of the *shi^549Δ3^* mutation, corresponding to the human dynamin-2-555Δ3 variant. *In vivo* models carrying this disease-associated deletion have not been reported to date. This approach enables analysis of the mutation in its endogenous genomic context, avoiding potential artifacts associated with overexpression-based systems. We examined disease-relevant phenotypes, including lethality, developmental defects, and impaired motility, and analyzed synaptic morphology and microtubule organization at larval neuromuscular junctions (NMJs). To explore the underlying pathogenic mechanisms, we performed a genetic interaction screen and identified *Patronin* as a modifier of *shi^549Δ3^* phenotypes*. Patronin* is the *Drosophila* homolog of calmodulin-regulated spectrin-associated proteins (CAMSAPs), which bind and stabilize microtubule minus-ends. We further investigated the functional interaction between dynamin and Patronin/CAMSAP *in vivo* and in cultured cells. Our findings provide mechanistic insights into how altered microtubule regulation contributes to neuronal defects associated with CMT-linked dynamin mutations.

## RESULTS

### *shi^549Δ3^* mutant flies exhibited developmental defects and shortened lifespan

First, we determined whether the dynamin-2-555Δ3 mutation causes a CMT disease-like phenotype in *Drosophila*. Using CRISPR/Cas9-mediated genome editing with a single-strand oligodeoxynucleotide donor, we generated a fly line carrying a precise nine-base deletion (*shi-549Δ3)*, which corresponds to the human dynamin-2-555Δ3 mutation (Figure 1A) ^21^. We examined the effects of *shi^549Δ3^* on development. Because *shi* is located on the X chromosome, male flies carrying the *shi^549Δ3^* mutation are hemizygous. These larvae exhibited a stepwise decline in the second- and third-instar stages compared with controls, suggesting that the *shi^549Δ3^* mutation induced developmental defects during the larval stages (Figure 1B). Consistently, the eclosion rate of hemizygous *shi^549Δ3^* mutant males was markedly decreased to approximately 3%, compared with 97% in FM7C males derived from the same cross (Figure 1C). In addition, the time from egg laying (crossing) to adult eclosion was delayed in both heterozygous and hemizygous *shi^549Δ3^* mutant flies (Figure S1).

**Figure 1.**
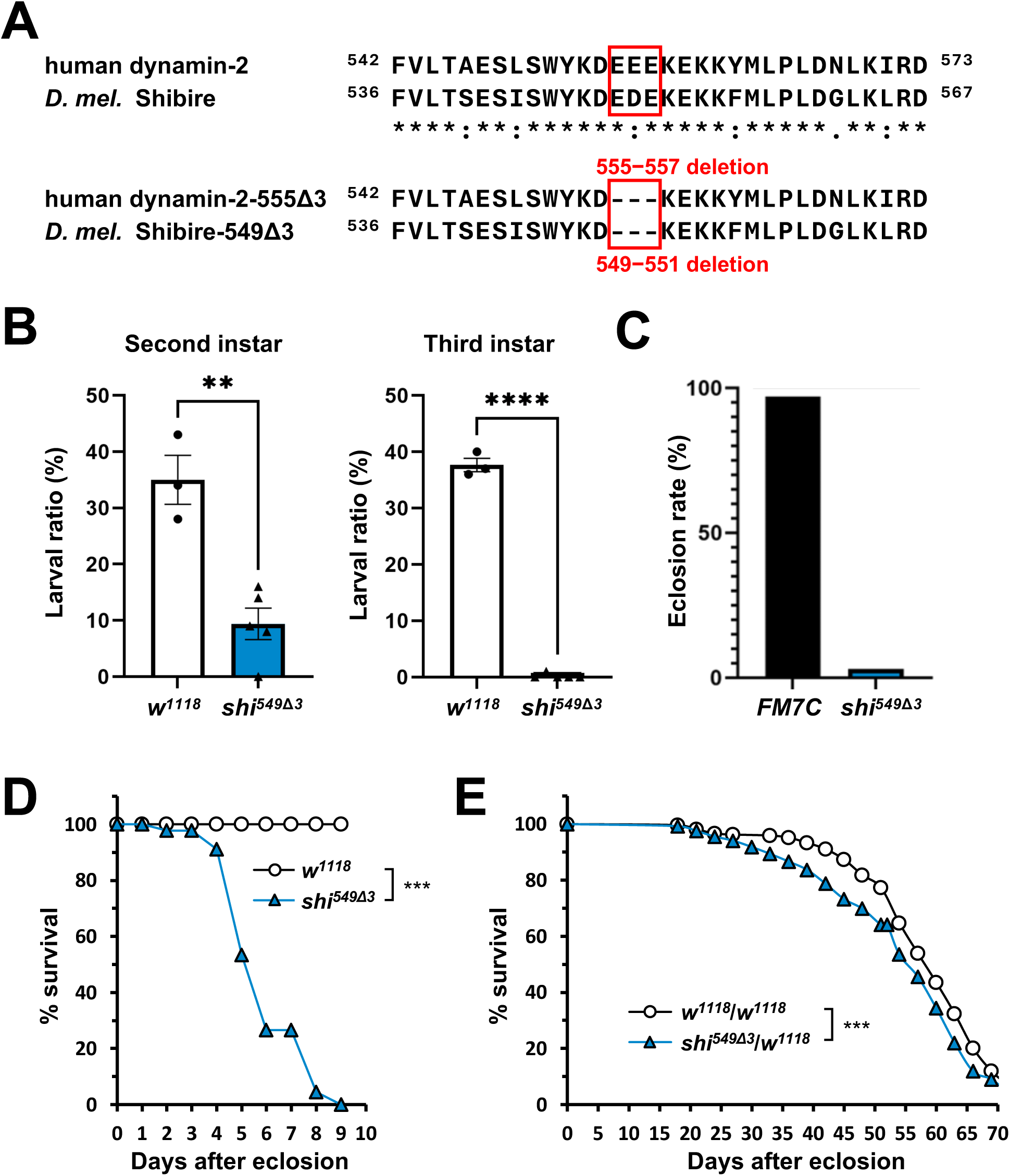
*shi*^549Δ3^ mutant flies exhibit developmental defects and shortened lifespans. (A) Alignment of amino acid sequences of human dynamin-2 (amino acids 542–573) and *Drosophila melanogaster (D. mel.)* Shibire (amino acids 536–567), performed using Clustal Omega (UniPlot). The corresponding mutant sequences are shown below the wild-type alignment. (B) Quantification of larval ratio (%) of control (*w^1118^*) and hemizygous *shi^549Δ3^* mutant larvae per vial. Flies were generated by crossing control *w^1118^*/FM7C, Tb females or *shi^549Δ3^*/FM7C, Tb females with FM7C, Tb males. The larval ratio was calculated as the percentage of non-Tubby larvae (corresponding to control or hemizygous mutant larvae) relative to total progeny (non-Tubby + Tubby). Each bar represents the mean ± s.e.m. of biologically independent experiments (control, n = 3; mutant, n = 5). Statistical analysis was performed using an unpaired t-test (**P < 0.01, ****P < 0.0001). (C) Percentage of control (FM7C) and hemizygous *shi^549Δ3^* mutant adult males among total adult male progeny. Flies were generated by crossing *shi^549Δ3^*/FM7C females with FM7C males. Male progeny were classified as FM7C males (control) or hemizygous *shi549Δ3* mutants (lacking FM7C). FM7C males accounted for 97% (109/112), whereas s*hi^549Δ3^* mutant males accounted for 3% (3/112). (D) Survival analysis of control (*w^1118^*) and hemizygous *shi^549Δ3^* mutant adult flies. Survival is expressed as the proportion of surviving individuals relative to day 0. Statistical significance was assessed using a log-rank test (control, n = 50; mutant, n = 45; ***P < 0.001). (E) Survival analysis of control (*w^1118^*/*w^1118^*) and heterozygous *shi^549Δ3^* mutant (*shi^549Δ3^*/*w^1118^*) adult flies. Survival is expressed as the proportion of surviving individuals relative to day 0. Statistical significance was assessed using a log-rank test (control, n = 279; mutant, n = 287; ***P < 0.001).

We next examined the effects of the *shi^549Δ3^* mutation on lifespan. The lifespans of both hemizygous and heterozygous adult mutants were significantly shortened compared with controls. Especially hemizygous mutants showed an exceptionally severe phenotype, as most individuals died within 9 days after eclosion (Figures 1D–1E). Collectively, these results demonstrate that the *shi^549Δ3^* mutation compromises development and survival in *Drosophila*.

### *shi^549Δ3^* mutant flies exhibited progressive motor dysfunction

Patients with CMT typically exhibit slowly progressive distal-dominant muscle weakness and sensory loss, which are attributed to damaged myelin and axons ^4,22^. Similarly, in *Drosophila*, impaired motor performance accompanied by neural defects, including morphological abnormalities at NMJs, have been reported in multiple CMT model flies ^23^. We therefore examined motor performance in mutant *shi^549Δ3^* flies to determine whether the *shi^549Δ3^* mutation induces CMT-like phenotypes.

Larval locomotor activity was assessed by a crawling assay on a grid (Figure 2A), which revealed that hemizygous *shi^549Δ3^* mutant third-instar larvae exhibited mildly impaired crawling ability compared with controls (Figure 2B). To further assess motor performance in adult flies, we conducted a forced climbing assay (Figure 2C). This assay demonstrated that both hemizygous and homozygous mutant *shi^549Δ3^* adults exhibited extremely severe climbing defects (Figures 2D–2E and S2A–S2B). By day 4 after eclosion, most hemizygous and homozygous mutant *shi^549Δ3^* flies were unable to walk and remained at the bottom of the vial. These results indicate that *shi^549Δ3^* mutants display more severe motor impairments during the adult stage than during the larval stage. By contrast, heterozygous *shi^549Δ3^* mutant flies exhibited a slow, age-dependent decline in climbing ability compared with controls (Figures 2F and S2C). The climbing defect in heterozygous *shi^549Δ3^* mutants became apparent around day 15, with mutants exhibiting decreased climbing ability compared with controls. However, by day 35, this difference was no longer evident due to an age-dependent decline in climbing ability in both mutant and control flies.

**Figure 2.**
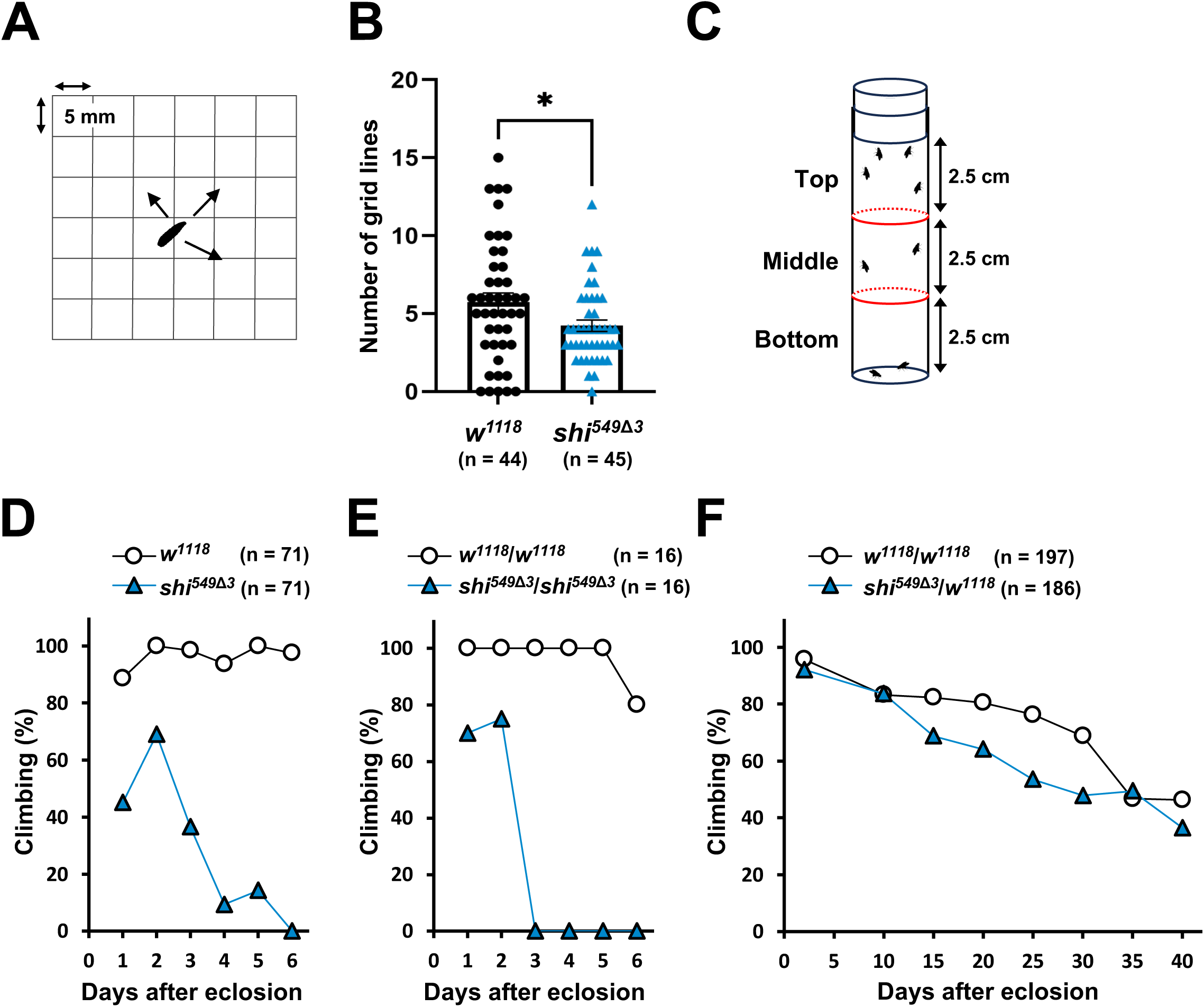
s*hi*^549Δ3^ mutant flies exhibit impaired locomotor activity. (A) Larval crawling was assessed on a 5 mm grid. Locomotor activity was quantified as the number of grid lines crossed by the posterior end of the larva within 2 min. (B) Crawling assay of wandering third-instar larvae. Locomotor activity is shown as the number of grid lines crossed by the posterior end of control (*w^1118^*, n = 44) and hemizygous *shi^549Δ3^* mutant (*shi^549Δ3^*, n = 45) larvae. Statistical analysis was performed using an unpaired t-test (*P < 0.05). Error bars indicate the mean ± s.e.m. (C) Adult climbing ability was quantified in vertical plastic vials (25 mm in diameter, 90 mm in height, marked with 25 mm intervals) by scoring the position of each fly 10 s after gently tapping the flies to the bottom of the vial. (D–F) Forced climbing assay of adult flies: control (*w^1118^* or *w^1118^/w^1118^*) and *shi^549Δ3^* mutants, including hemizygous (*shi^549Δ3^*, D), homozygous (*shi^549Δ3^*/*shi^549Δ3^*, E), and heterozygous (*shi^549Δ3^*/*w^1118^*, F) animals. Results are shown as the percentage of flies that reached the top of the vial. See also Figure S2 for the detailed distribution of flies across each region: hemizygous *shi^549Δ3^* males (Figure S2A), homozygous *shi^549Δ3^*/*shi^549Δ3^* females (Figure S2B), and heterozygous *shi^549Δ3^*/*w^1118^* females (Figure S2C).

Altogether, the observed progressive climbing impairment (Figure 2F), delayed eclosion (Figure S1), and shortened lifespan (Figure 1E) indicate that heterozygous mutant *shi^549Δ3^* flies exhibit measurable functional deficits. These results suggest that mutant *shi^549Δ3^* flies exhibit early-onset and slowly progressive motor dysfunction, a feature reminiscent of the disease course observed in many patients with CMT ^23^.

### The *shi^549Δ3^* mutation induced synaptic overgrowth at NMJs

The NMJ is a common target in neuromuscular and peripheral nerve disorders. NMJ dysfunction has been reported in many CMT models and is suggested to contribute to the pathogenesis of CMT ^24,25^. Therefore, to investigate NMJ morphology in mutant *shi^549Δ3^* flies, hemizygous mutant larval fillets were stained with antibodies against horseradish peroxidase (HRP), a neuronal membrane marker, and Discs large, a primarily postsynaptic scaffold protein (Figure 3A). Mutant *shi^549Δ3^* flies showed increased axonal branching (Figures 3B and S3A) and a higher number of type Ib boutons (large glutamatergic synaptic boutons) ^26^ compared with controls (Figures 3C and S3B–C). In addition, the *shi^549Δ3^* mutation induced the formation of small ectopic satellite boutons, a marker of excessive bouton overgrowth ^27,28^ (Figure 3A, white arrows; Figures 3D and S3D). By contrast, the number of ghost boutons, immature boutons transiently formed during bouton development ^29^, was not increased in *shi^549Δ3^* mutant flies (Figures 3E and S3E). These results indicate that the *shi^549Δ3^* mutation promotes synaptic overgrowth at NMJs.

**Figure 3.**
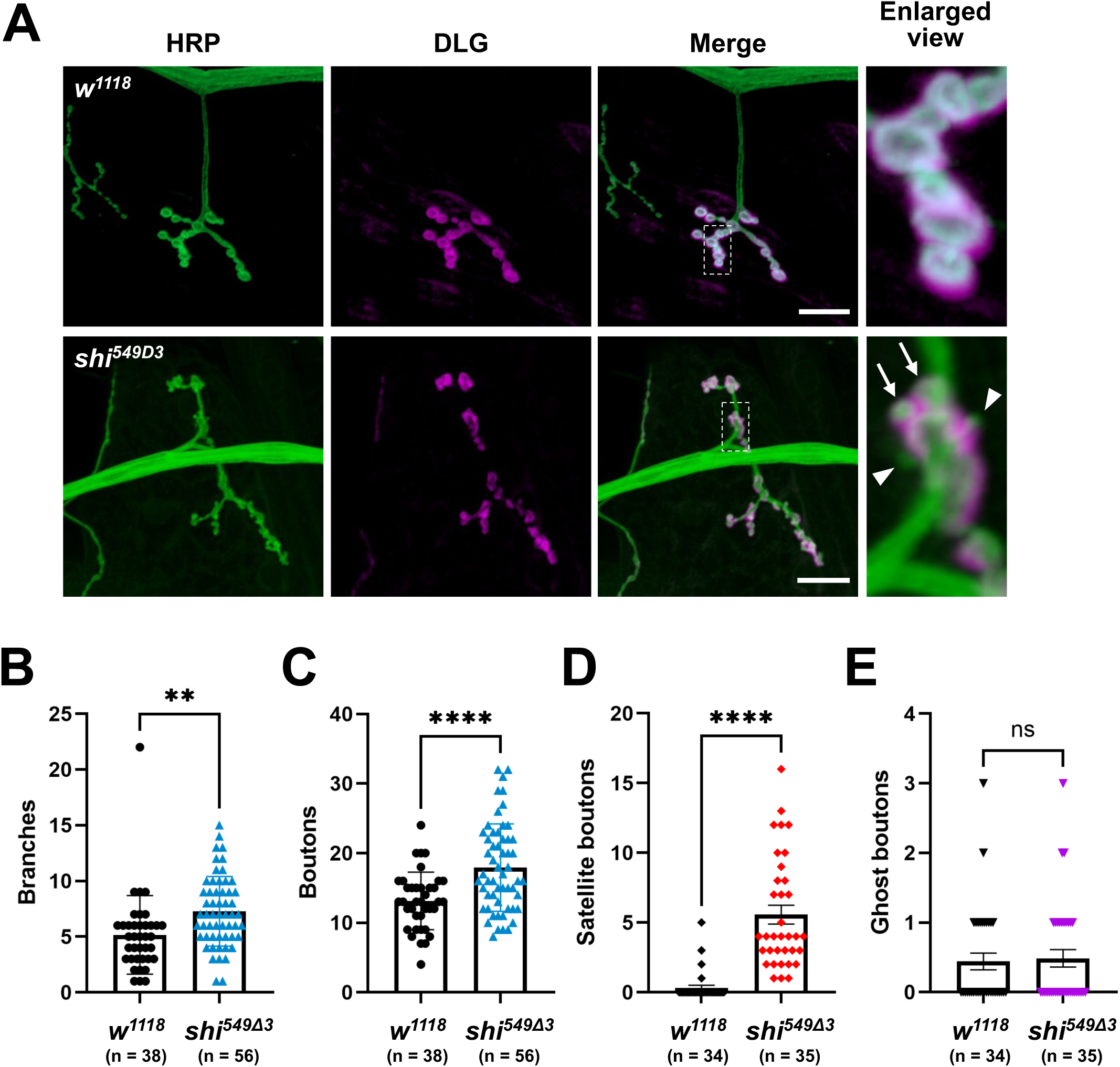
Altered synaptic bouton morphology at the neuromuscular junctions (NMJs) of *shi^549Δ3^* mutant larvae. (A) Representative images of NMJs at muscle 4 in wandering third-instar larvae stained with antibodies against HRP (green) and DLG (magenta). All genotypes are hemizygous control (*w^1118^*) or *shi^549Δ3^* mutant larvae. Hemizygous larvae were identified as non-Tubby progeny based on the absence of the Tb marker. Scale bars, 20 μm. White frames indicate magnified views of terminal boutons containing satellite boutons (arrows) and ghost boutons (arrowheads). (B–E) Quantification of the numbers of (B) axonal branches, (C) type Ib boutons, (D) satellite boutons, and (E) ghost boutons at NMJs of muscle 4 in abdominal segments A1– A4. Details of the bouton numbers in each individual A1–A4 muscle segment are shown in Figure S3C. For (B, C), NMJs (control, n = 38 from 5 larvae; mutant, n = 56 from 11 larvae) were analyzed. For (D, E), NMJs (control, n = 34 from 5 larvae; mutant, n = 35 from 11 larvae) were analyzed. Statistical analysis was performed using an unpaired t-test (*P < 0.05, **P < 0.01, ****P < 0.0001; ns, no significance). Error bars indicate the mean ± s.e.m.

### The *shi^549Δ3^* mutation induced excessive microtubule stabilization at NMJs

Previous studies have shown that dynamin-2-555Δ3 and its splicing isoform dynamin-2*-* 551Δ3 strongly colocalize with microtubules and promote their acetylation and stabilization in mammalian cultured cells ^19,20^. Increased levels of stable or dynamic microtubules can trigger excessive formation of synaptic boutons, including the budding of satellite boutons ^30–32^. Therefore, we investigated whether microtubule stability is increased in *shi^549Δ3^* mutant flies. First, α-tubulin staining was performed to visualize alterations in the localization and extension of microtubules at larval NMJs (Figure 4A). The number of terminal boutons containing bundled microtubules was increased in the *shi^549Δ3^* mutants compared with the controls, suggesting that microtubules extend further into the terminal boutons (Figures 4A–4B). Similarly, acetylated α-tubulin, a marker of stable microtubules ^33^, was also observed to extend into the terminal boutons in the *shi^549Δ3^* mutants (Figures 4C–4D).

**Figure 4.**
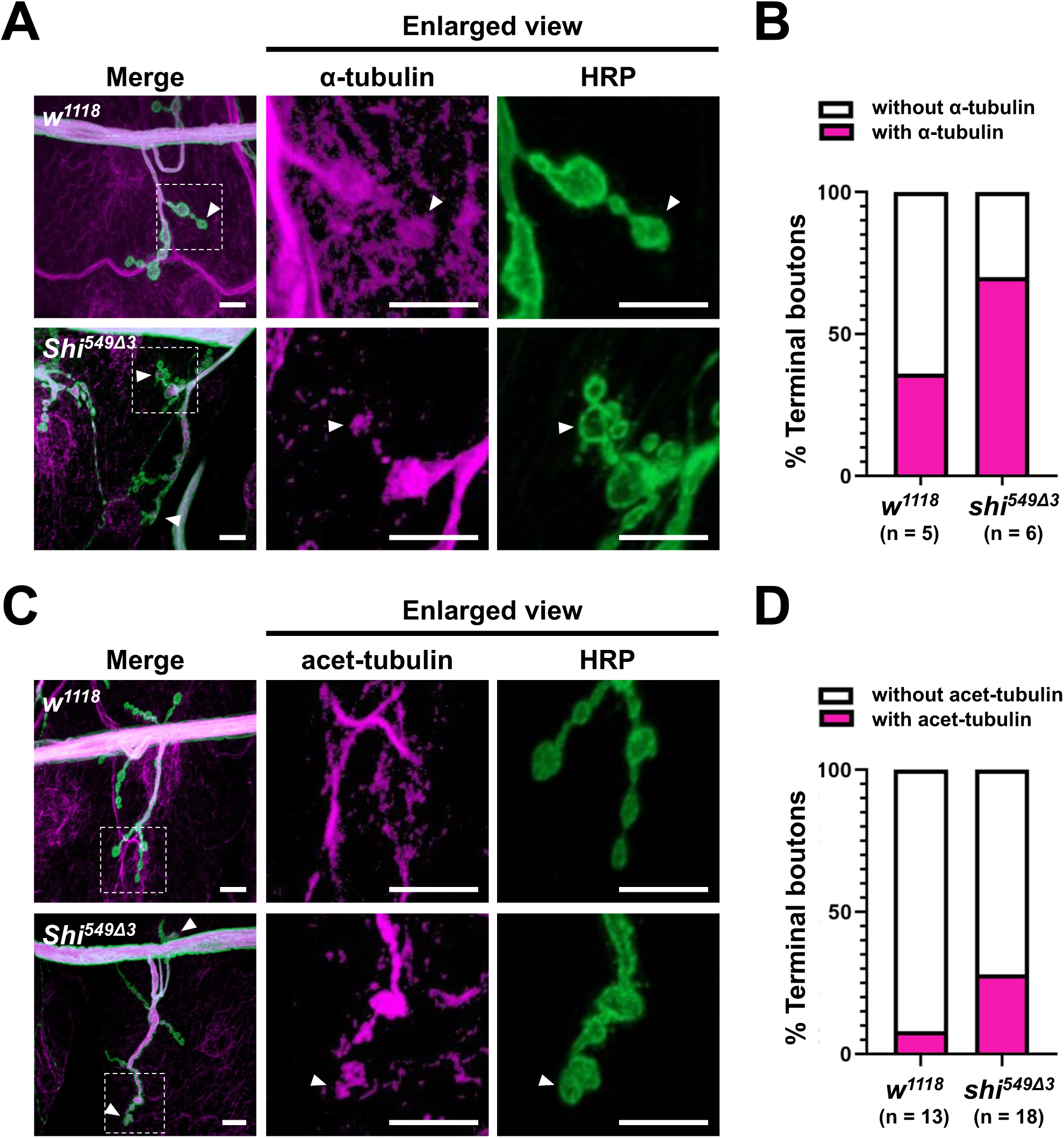
The *shi*^549Δ3^ mutation induces excessive microtubule stabilization at NMJs. (A, C) Representative images of larval NMJs from control (*w^1118^*) and hemizygous *shi^549Δ3^* mutant larvae stained with antibodies against α-tubulin (A, magenta), acetylated α-tubulin (acet-tubulin; C, magenta), and HRP (green). Arrowheads indicate terminal boutons containing bundled or acetylated α-tubulin. Scale bars, 10 μm. (B, D) Quantification of the proportion of terminal boutons containing (B) bundled α-tubulin or (D) acetylated α-tubulin relative to the total number of terminal boutons at NMJs on muscle 4 in abdominal segments A1–A4. For (B), NMJs (control, n = 5 from 3 larvae; mutant, n = 6 from 1 larva) were analyzed. For (D), NMJs (control, n = 13 from 5 larvae; mutant, n = 18 from 3 larvae) were analyzed.

We further examined the effects of the *shi^549Δ3^* mutation on microtubule stability by staining for Futsch, which is homologous to the mammalian microtubule-associated protein 1B ^34,35^. Futsch colocalizes with stable microtubules and forms loop structures (“Futsch loops”) in boutons ^36,37^. To quantify Futsch distribution, Futsch staining patterns in boutons were classified into four categories: looped, continuous (bundles or splayed bundles), diffuse, and negative (undetectable) based on a previous report ^38^ (Figure S4A). Satellite boutons were excluded from the analysis, and the numbers of boutons per NMJ were quantified. The *shi^549Δ3^* mutants showed an increased number of boutons containing Futsch loops (Figure S4B), continuous Futsch signals (Figure S4C), and diffuse Futsch signals (Figure S4D). By contrast, the numbers of boutons lacking detectable Futsch signals were unchanged (Figure S4E). These results suggest that *shi^549Δ3^* promotes microtubule stabilization in synaptic terminals.

To test this possibility, we used a pharmacological approach. Control and *shi^549Δ3^* larval fillets were treated with nocodazole, a microtubule-destabilizing agent ^39^ (Figures 5A and 5D). Low concentrations of nocodazole decreased the number of Futsch loops in *shi^549Δ3^* larvae (Figures 5D and 5E) but had no effect in control larvae (Figures 5A and 5B), consistent with a previous report ^32^. By contrast, treatment with paclitaxel, a microtubule-stabilizing agent ^40,41^, increased the number of Futsch loops in control larvae (Figures 5A and 5B) but had no effect in *shi^549Δ3^* larvae (Figures 5D and 5E). Neither nocodazole nor paclitaxel affected the numbers of type Ib boutons (Figures 5C and 5F), consistent with a previous report ^32^. Most likely, the exposure time was too short to affect synaptic morphogenesis. Nevertheless, these results support the idea that the *shi^549Δ3^* mutation causes excessive stabilization of microtubules at synaptic terminals, thereby promoting excessive synaptic growth.

**Figure 5.**
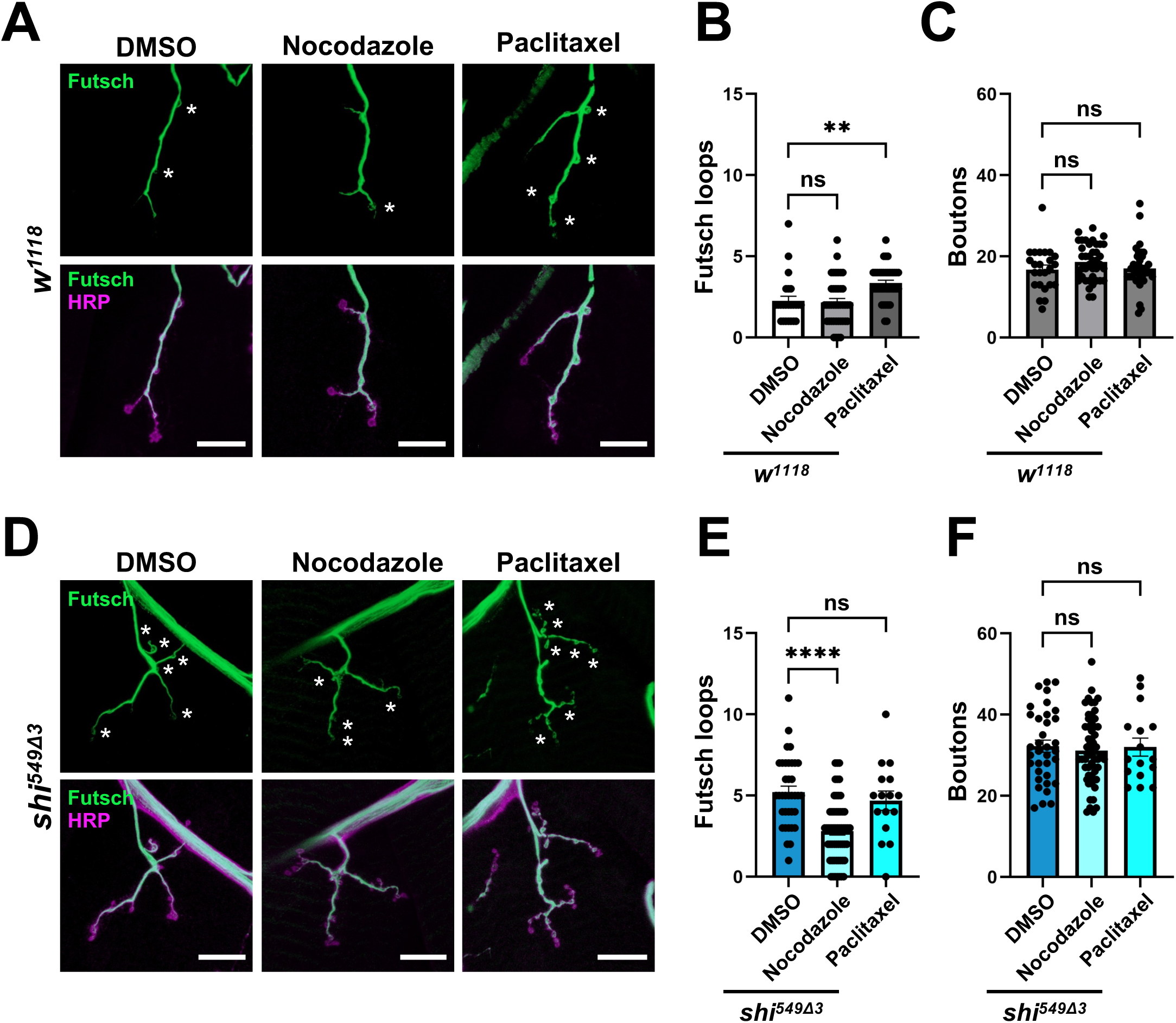
Nocodazole suppresses Futsch-positive microtubule loop formation at NMJs in *shi*^549Δ3^ mutants. (A, D) Representative images of larval NMJs from (A) control (*w^1118^*) and (D) hemizygous *shi^549Δ3^* mutant larvae stained with antibodies against Futsch (green) and HRP (magenta). Larval fillets were treated with DMSO (vehicle control; 30 min or 1 h), 50 μM paclitaxel (30 min), or 30 μM nocodazole (1 h) prior to fixation. Asterisks indicate Futsch-positive microtubule loops (“Futsch loops”) in type Ib boutons. Scale bars, 20 μm. (B, E) Quantification of the number of Futsch loops at NMJs on muscle 4 in abdominal segments A1–A4 from control (B) and hemizygous *shi^549Δ3^* mutant (E) larvae. (C, F) Quantification of the number of type Ib boutons at the same NMJs from control (C) and hemizygous *shi^549Δ3^* mutant (F) larvae. For DMSO treatment, NMJs (control, n = 24 from 4 larvae; mutant, n = 36 from 6 larvae) were analyzed. For nocodazole treatment, NMJs (control, n = 40 from 9 larvae; mutant, n = 56 from 8 larvae) were analyzed. For paclitaxel treatment, NMJs (control, n = 37 from 8 larvae; mutant, n = 16 from 4 larvae) were analyzed. Statistical analysis was performed using Dunnett’s multiple comparisons test (**P < 0.01, ****P < 0.0001; ns, not significant). Error bars indicate the mean ± s.e.m.

### Genetic depletion of Patronin rescued peripheral neuropathy-like phenotypes in *shi^549Δ3^* mutants

To elucidate the molecular mechanisms underlying the excessive stabilization of microtubules induced by the *shi^549Δ3^* mutation, we performed genetic screening to identify factors that interact with *shi^549Δ3^*. Heterozygous female *shi^549Δ3^* mutants were crossed with males carrying deficiencies or mutations in candidate genes, and modifiers that rescued developmental defects were screened. *Patronin* was identified as a genetic interactor of *shi^549Δ3^*. *Drosophila* Patronin is homologous to the mammalian CAMSAP family ^42^. The Patronin/CAMSAP family comprises conserved proteins that bind to microtubule minus-ends. These proteins are required for the formation of non-centrosomal microtubules and protect microtubule minus-ends from kinesin-13-dependent depolymerization ^43–46^.

Two P-element insertion lines (*Patronin^k07433^, Patronin^EY05252^* and *P{GSV7}GS21406*) ^47^ increased the eclosion rate of *shi^549Δ3^* mutants from 3% to 21%, 29%, and 15%, respectively (Figure 6A, red region). Eclosion rescue was also confirmed by pan-neuronal and motor neuron-specific knockdown of *Patronin* using *elav-Gal4* and *C164-Gal4*, respectively (Figure 6A, green region). Furthermore, pan-neuronal and motor neuron-specific knockdown of *Patronin* almost completely rescued the climbing ability of *shi^549Δ3^* mutants (Figure 6B). These results indicate that the peripheral neuropathy-like phenotypes of *shi^549Δ3^* mutants depend, at least in part, on Patronin function, suggesting that dysregulation of microtubule minus-end stability contributes to their pathogenesis.

**Figure 6.**
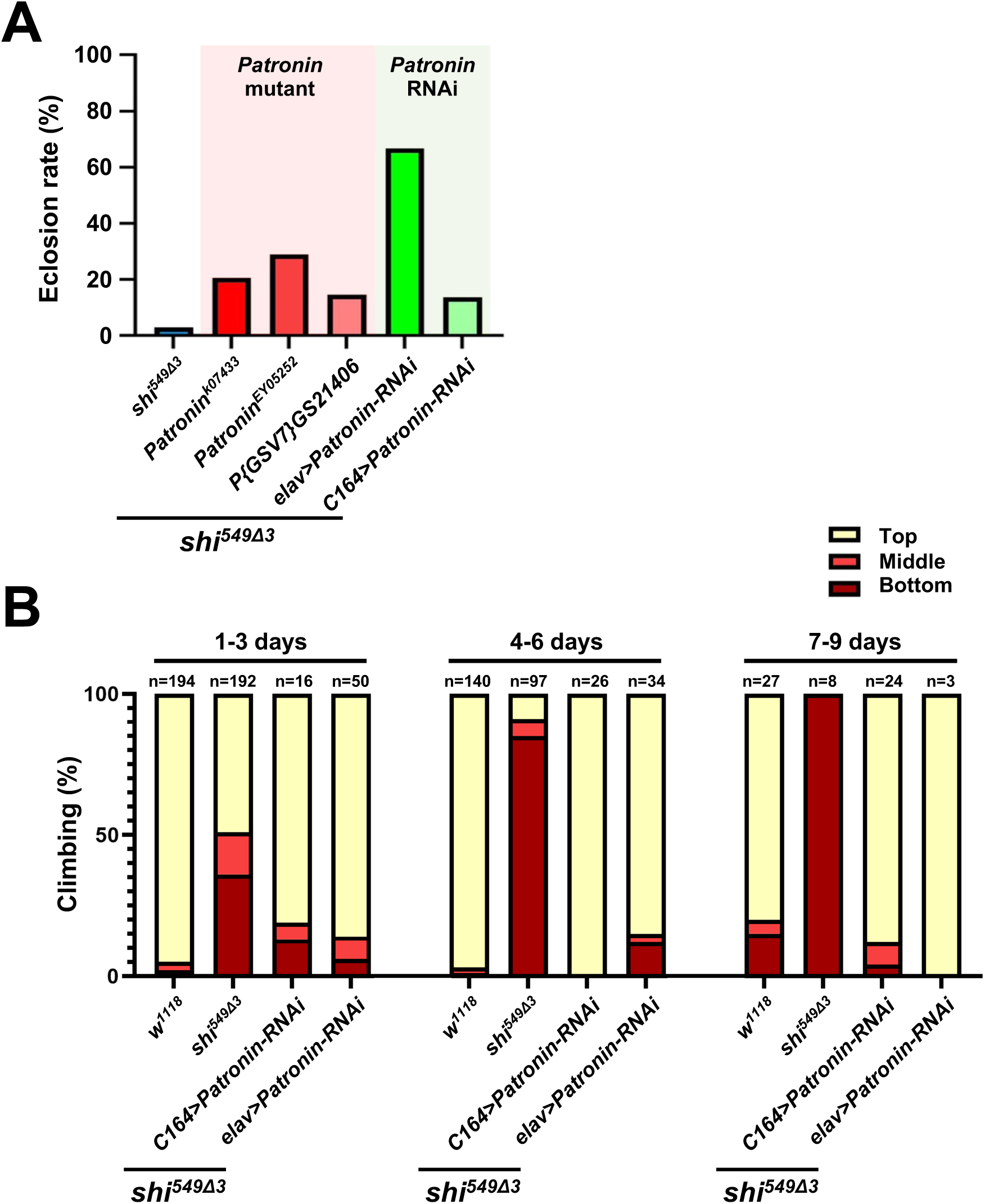
Genetic depletion of *Patronin* rescues lethality and improves locomotor performance in *shi*^549Δ3^ mutants. (A) Percentage of hemizygous *shi^549Δ3^* mutant adult males carrying P-element insertion *Patronin* alleles (*Pat^K^*^07433^, *Pat^EY^*^05252^, and *P{GSV7}GS21406*; red) or neuron-specific *Patronin* knockdown (*elav-Gal4 or C164-Gal4 > UAS-Patronin-RNAi*; green), relative to the total number of adult males in the progeny. Hemizygous *shi^549Δ3^* adult males were used as controls. The percentages (number/total) are as follows: *shi^549Δ3^*, 3% (3/112); *shi^549Δ3^*; *Patronin^K^*^07433^, 21% (15/73); *shi^549Δ3^*; *Patronin^EY^*^05252^, 29% (20/69); *shi^549Δ3^*; *P{GSV7}GS21406*, 15% (7/48); *shi^549Δ3^*; *elav-Gal4 > UAS-Patronin-RNAi*, 68% (19/28); *shi^549Δ3^*; *C164-Gal4 > UAS-Patronin-RNAi*, 14% (17/125). (B) Forced climbing assay of control (*w^1118^*) and hemizygous *shi^549Δ3^* mutant adult males, including animals with neuron-specific *Patronin* knockdown (*elav-Gal4 or C164-Gal4 > UAS-Patronin-RNAi*). Results are presented as the percentage of flies in the top (yellow), middle (red), or bottom (dark red) regions of the vials. The number above each graph indicates the age of the flies (days post-eclosion). Sample sizes for each dataset are indicated in the graphs.

### CAMSAP2 knockdown suppressed dynamin-2-induced microtubule over-acetylation in HeLa cells

To test whether the effect observed in *Drosophila* is conserved in mammalian cells, and whether CAMSAP2 mediates dynamin-2-555Δ3-induced excessive microtubule acetylation ^19,20^, we performed CAMSAP2 knockdown using shRNA in HeLa cells (Figure 7A). In control HeLa cells, CAMSAP2 knockdown significantly reduced microtubule acetylation (Figures 7A–7B). Moreover, CAMSAP2 knockdown reduced microtubule acetylation to control levels in HeLa cells expressing hemagglutinin (HA)- tagged dynamin-2-555Δ3 (Figures 7A–7B). These results demonstrate that CAMSAP2 is required for dynamin-2-induced microtubule over-acetylation and support a conserved mechanism by which dynamin regulates microtubule dynamics across various animal species, from *Drosophila* to mammals.

**Figure 7.**
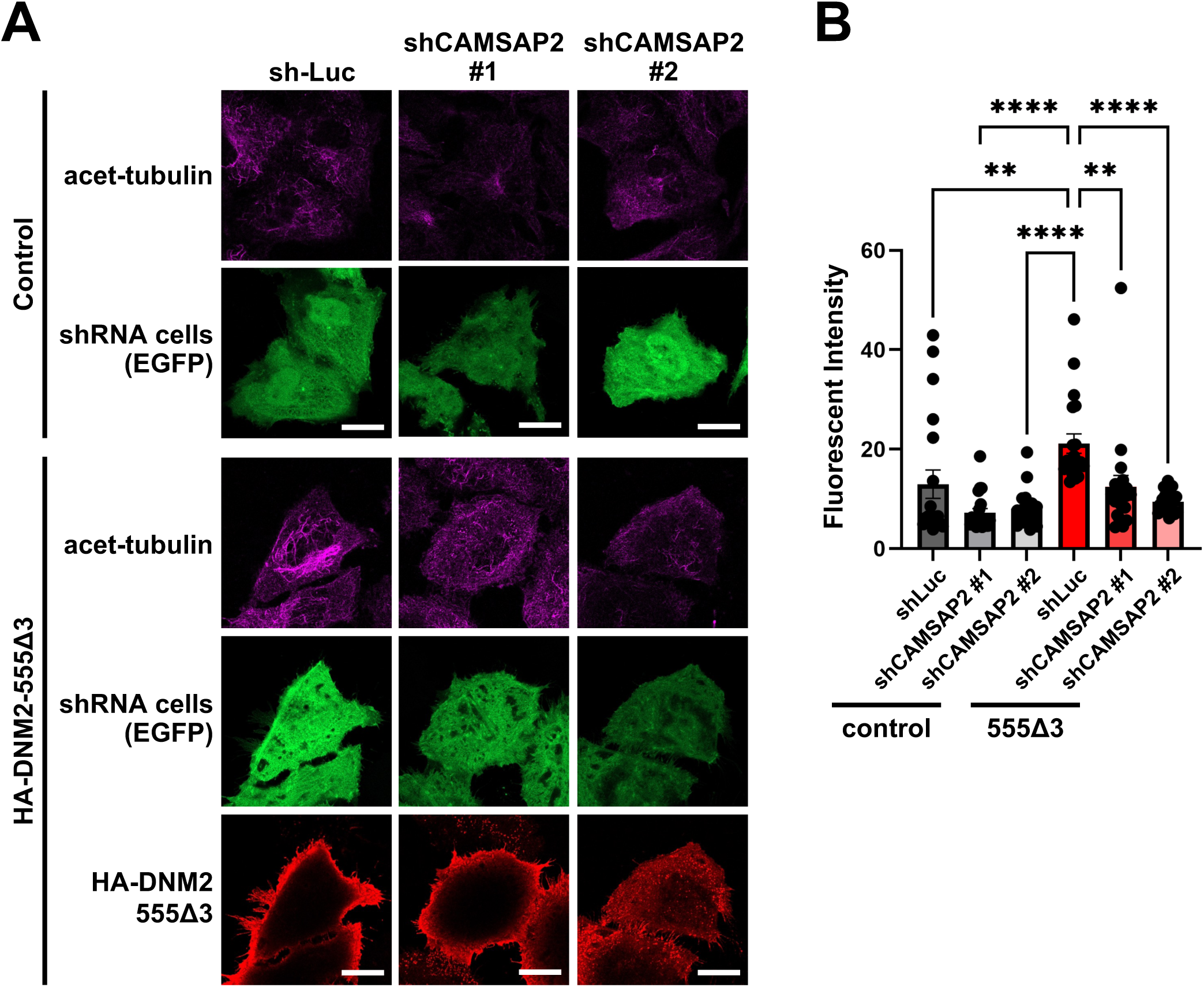
CAMSAP2 depletion suppresses DNM2-555Δ3-induced microtubule acetylation. (A) Representative images of HeLa cells expressing or not expressing HA-DNM2-555Δ3 together with shLuciferase (shLuc) or shCAMSAP2 (#1 or #2). Cells expressing shRNAs are identified by co-expression of EGFP (green). Cells were stained with antibodies against acetylated α-tubulin (acet-tubulin, magenta) and HA (red). Scale bars, 20 μm. (B) Quantification of total acetylated α-tubulin fluorescence intensity per cell for the experiments shown in (A). Statistical analysis was performed using Dunnett’s multiple comparisons test (**P < 0.01, ****P < 0.0001; n = 20 cells per condition). Error bars indicate the mean ± s.e.m.

## DISCUSSION

Here we show that a *Drosophila* model carrying the CMT-associated *shi^549Δ3^* mutation recapitulates key features of DI-CMTB, thereby providing insight into the cellular mechanisms underlying axonal pathology. Heterozygous *shi^549Δ3^* mutant flies exhibited slowly progressive motor impairment (Figure 2F), resembling that observed in patients with DI-CMTB. In addition, abnormal axonal morphology at synaptic terminals in hemizygous *shi^549Δ3^* mutant larvae (Figure 3) is consistent with axonal abnormalities characteristic of DI-CMTB ^6,48^. Together, these findings indicate that *shi^549Δ3^* mutant flies reproduce several hallmarks of CMT-related pathology and may thus serve as a useful model.

We also found that the *shi^549Δ3^* mutation increased microtubule stability in *Drosophila* (Figures 4 and S4), consistent with previous observations in mammalian cultured cells ^19,20^. At *Drosophila* NMJs, microtubule stability (often assessed by Futsch loops formation) is correlated with bouton number ^31,32^. Consistently, *shi^549Δ3^* mutant larvae showed enlarged Futsch loops together with increased bouton numbers, suggesting that microtubule stabilization contributes to bouton overgrowth. However, this relationship is not always straightforward. Several studies have shown that genetic manipulations affecting microtubule acetylation, such as mutation of the K394 acetylation site of α-tubulin ^32^ or deletion of the *Drosophila* α-tubulin acetyltransferase dαTAT ^31^, can also increase bouton number despite decreasing or not affecting Futsch loops. These findings suggest that bouton growth is regulated by multiple factors and pathways, including actin dynamics and bone morphogenetic protein signaling ^37,49^, and that microtubule stabilization alone cannot explain all aspects of bouton overgrowth. In addition, endocytosis-defective mutants, such as temperature-sensitive *shi^TS^* strains, exhibit increased bouton numbers, often accompanied by satellite boutons, similar to those observed in *shi^549Δ3^* mutants ^50–52^. This phenotype has been interpreted as a compensatory response to reduced synaptic transmission that includes an enhanced active zone and increased T-bar density. Future studies should investigate whether *shi^549Δ3^* mutants also exhibit endocytic defects, as impaired dynamin function may affect both endocytosis and microtubule dynamics, and the relative contributions of these processes to synaptic overgrowth remain to be clarified.

The dynamin-2-555Δ3 mutation has been reported to act in a gain-of-function manner, with enhanced self-assembly activity ^53^. Consistent with this, immunostaining revealed that dynamin-2-555Δ3 localizes more strongly to microtubules compared with wild-type dynamin-2 ^19^, suggesting an increased interaction with microtubules. It is possible that the 555Δ3 mutation enhances abnormal interactions with microtubules, thereby shifting the balance of microtubule dynamics toward excessive stabilization.

Patronin/CAMSAPs stabilize microtubules by binding to their minus-ends. In *Drosophila*, decreased Patronin expression partially improved both lethality and climbing defects in *shi^549Δ3^* mutants, suggesting that excessive microtubule stabilization contributes, at least in part, to the observed phenotypes. Consistent with this idea, CAMSAP2 knockdown in dynamin-2-555Δ3-overexpressing HeLa cells decreased acetylated tubulin levels, supporting a role for Patronin/CAMSAP in promoting microtubule stability in our experimental system. It is possible that aberrant dynamin activity enhances microtubule stabilization, at least in part, through a Patronin/CAMSAP-dependent mechanism. Although the precise molecular relationship between dynamin and Patronin/CAMSAP remains to be determined, our findings suggest that regulation of microtubule minus-end dynamics may represent a key pathway downstream of dynamin.

Taken together, our findings demonstrate that the disease-associated *shi^549Δ3^* mutation induces peripheral neuropathy-like phenotypes accompanied by excessive microtubule stabilization at synaptic terminals. We identify dysregulated microtubule stabilization as a key neuropathogenic mechanism downstream of dynamin dysfunction that is modulated by the microtubule minus-end regulator Patronin/CAMSAP. These results establish the *Drosophila shi^549Δ3^* model as a tractable *in vivo* system and provide mechanistic insight into dynamin-related neuropathies.

## ACKNOWLEDGMENTS

We would like to thank Ms. F. Terada (Hiroshima University, Japan) for generating *shi^549Δ3^* mutant flies. We are also grateful to Bloomington Stock Center, Vienna Drosophila Resource Center, and KYOTO Drosophila Stock Center for the fly stocks used in this study. We also thank Enago (www.enago.jp) for English language editing of the manuscript.

This work was supported by JSPS KAKENHI (Grant Numbers JP23K06483 to KH and JP24K02062 to TC). This work was also supported by grants from Narishige Zoological Science Award and Tsuchiya Memorial Medical Foundation to KH.

## AUTHOR CONTRIBUTIONS

K.H. conceived the study. Y.M. performed the experiments and analyzed the data. Y.M., M.O., T.C., and K.H. designed the experiments and wrote the manuscript.

## MATERIALS AND METHODS

### Fly strains

The following *D. melanogaster* strains were used: *w^1118^*, *elav-Gal4, C164-Gal4, ubi-Gal4, Patronin^k^*^07433^ (BL#10672), and *Patronin^EY^*^05252^ (BL#16647) (Bloomington *Drosophila* Stock Center, Indiana University, USA), *P{GSV7}GS21406* (#202731) (*Drosophila* Genetic Resource Center, Kyoto, Japan), *Patronin-RNAi* (v108927) (Vienna *Drosophila* RNAi Center), and *shi^549Δ3^* (this study). Flies were reared on standard fly food at 25°C under a 12 h light/12 h dark cycle with approximately 60% relative humidity.

### Generation of the *shi^549Δ3^* fly strain

To generate the *shi^549Δ3^* strain, sense and antisense oligonucleotides were designed using flyCRISPR to construct a gRNA expression plasmid. Annealed oligonucleotide pairs were ligated into the *U6b* plasmid digested with *Bbs*I using Ligation High Ver.2 (Toyobo, Japan). A single-strand oligodeoxynucleotide was used as a donor template for the generation of the *shi^549Δ3^* allele. The following oligonucleotides were used: gRNA target sense, CTTCGCGAGAGTATCTCCTGGTATA; gRNA target antisense, AAACTATACCAGGAGATACTCTCGC; and donor oligonucleotide, TTTAAACCATCCAAACCATCCAACGGCAGCATAAATTTCTTCTCCTTCTCTTTA TACCAGGAGATACTCTCGGATGTCAGCACAAACCAATACGGACGCGATCCA. The gRNA plasmid and single-strand oligodeoxynucleotide were injected into *Drosophila* embryos by BestGene Inc. (Chino Hills, CA, USA). Injected embryos developed to adulthood, and the resulting flies were crossed to balancer stocks. Progeny were then screened for the presence of the mutation by high-resolution melt analysis, and candidate flies were further validated by genomic DNA sequencing.

### Measurements of fly development

Flies (10 females and 5 males per vial) were maintained at 25°C with standard fly food and allowed to mate. To assess larval development, 4 days after mating, a 15% (w/v) sucrose solution was added to each vial at a volume equal to that of the food, and the food surface was gently stirred with a spatula to allow larvae to float to the surface. Floating larvae were collected, transferred to Petri dishes, and counted. Larvae were classified by genotype and developmental stage under a stereomicroscope. To assess eclosion, newly eclosed adults were collected daily, sorted by genotype, and counted.

### Lifespan assay

Approximately 10 newly eclosed female flies were housed per plastic vial containing 5– 10 ml of standard fly food. Flies were maintained at 25°C and 60% relative humidity under a 12 h light/12 h dark cycle. Flies were transferred to fresh food three times per week, and the number of dead flies was recorded. For extremely short-lived hemizygous *shi^549Δ3^* mutants, flies within 24 h after eclosion were transferred to fresh vials containing standard fly food and maintained at 25°C and 60% relative humidity under a 12 h light/12 h dark cycle. Survival was recorded every 24 h.

### Larval locomotor assay

Fifteen third-instar larvae were washed with phosphate-buffered saline (PBS) and transferred onto a 1% agarose gel in an 80 mm Petri dish marked with grid lines at 5 mm intervals. Larvae were allowed to acclimatize for 1 min, and locomotor activity was assessed by counting the number of grid lines crossed by the posterior end of each larva within 2 min. Assays were performed at room temperature (RT).

### Forced climbing assay

Flies within 24 h after eclosion were separated by sex and transferred to fresh vials containing standard fly food. Flies were maintained at 25°C and 60% relative humidity under a 12 h light/12 h dark cycle and transferred to fresh fly food every 2–3 days. Next, they were transferred to empty plastic vials (25 mm in diameter and 90 mm in height) marked with three horizontal lines at 25 mm intervals from the bottom and allowed to acclimatize for 1 min. Flies were gently tapped to the bottom of the vial, and the numbers of flies located at the bottom, middle, and top regions were recorded 10 s after tapping. After each measurement, flies were returned to their original food-containing vials. Measurements were repeated daily until all flies had died. The percentage of flies present at each height were calculated. Assays were performed at RT.

### Immunostaining and imaging of NMJs

Wandering third-instar larvae were dissected in PBS and fixed in 4% paraformaldehyde in PBS for 20 min. Samples were blocked with 5% normal goat serum in PBS containing 0.3% Triton-X100 (PBST) for at least 1 h at RT. Larval fillets were incubated with primary antibodies diluted in PBST overnight at 4°C, washed with PBST at RT, and incubated with secondary antibodies and tetramethylrhodamine-isothiocyanate-conjugated phalloidin (1:250, Sigma–Aldrich) diluted in PBST overnight at 4°C in the dark. After washing with PBST, samples were mounted in SlowFade (Thermo Fisher Scientific). The following primary antibodies were used: mouse anti-Discs large (1:25; Developmental Studies Hybridoma Bank) and rabbit anti-HRP (1:125; Jackson ImmunoResearch). The following secondary antibodies were used: goat anti-mouse IgG Alexa Fluor 633 and goat anti-rabbit IgG Alexa Fluor 488 (both 1:250; Thermo Fisher Scientific). Images were acquired using a confocal microscope (LSM900; Carl Zeiss AG, Oberkochen, Germany). Axon branch number and bouton number were quantified using Fiji (ImageJ, NIH, v.1.49k) with the macros “*Drosophila* NMJ Morphometrics” and “*Drosophila* NMJ Bouton Morphometrics” as described previously ^54^. Bouton number was quantified at NMJ4 in abdominal segments A1–A4 due to its relative simplicity.

### Immunostaining and imaging of tubulin markers at NMJs

Wandering third-instar larvae were dissected in BRB80 buffer (400 mM PIPES, pH 6.8, 25 mM EGTA, 5 mM MgCl2) and then fixed, blocked, stained, and mounted as described above. The following primary antibodies were used: mouse anti-Futsch (22C10, 1:50; Developmental Studies Hybridoma Bank), mouse anti-α-tubulin (clone DM1A, 1:500; CEDARLANE), mouse anti-acetylated tubulin (1:500; Merck, Darmstadt, Germany), and rabbit anti-HRP (1:125; Jackson ImmunoResearch). Tetramethylrhodamine-isothiocyanate-conjugated phalloidin (1:250; Sigma–Aldrich) and the following secondary antibodies were used: goat anti-mouse IgG Alexa Fluor 633, goat anti-mouse IgG Alexa Fluor 488, goat anti-rabbit IgG Alexa Fluor 488, and goat anti-rabbit IgG Alexa Fluor 680 (all 1:250; Thermo Fisher Scientific).

### Paclitaxel and nocodazole treatment

Prior to fixation, wandering third-instar larvae were dissected in BRB80 buffer containing either 50 μM paclitaxel (Sigma–Aldrich) or 30 μM nocodazole (Calbiochem) and incubated for 30 min or 1 h. BRB80 buffer containing 1% DMSO was used as a control. Tissues were then fixed and stained as described above.

### Cell culture and transfection

HeLa cells (RCB0007) were obtained from RIKEN BRC through the National Bio-Resource Project of MEXT/AMED, Japan. Cells were maintained in Eagle’s Minimum Essential Medium (Nissui Pharmaceuticals) supplemented with 10% (v/v) fetal bovine serum (Biological Industries) at 37°C in a humidified atmosphere containing 5% CO2.

The pcDNA3.1-*HA-DNM2-555Δ3* construct was generated from pcDNA3.1-*HA-DNM2* (kindly provided by Dr. S. L. Schmid) by PCR-based mutagenesis and used for the overexpression experiments. For CAMSAP2 knockdown, short hairpin RNAs targeting CAMSAP2 were designed and cloned into the pBluescript SK(+) vector under the control of the U6 promoter. Two independent shRNAs targeting CAMSAP2 (shCAMSAP#1 and shCAMSAP#2) were used, with a luciferase-targeting shRNA (shLuc) serving as a negative control. The target sequences were as follows: shCAMSAP#1, 5 ′ - GTACTGGATAAATAAGGTACTCGAGTACCTTATTTATCCAGTAC-3 ′ ; shCAMSAP#2, 5 ′ - GAATACTTCTTGACGAGTTCTCGAGAACTCGTCAAGAAGTATTC-3 ′ ; shLuc, 5 ′ - GCGTACGCGGAATACTTCGACTCGAGTCGAAGTATTCCGCGTAC-3 ′ . The vector also encodes EGFP under the control of the CMV promoter, allowing identification of shRNA-expressing cells. Cotransfection was performed using Lipofectamine LTX Reagent (Thermo Fisher Scientific) according to the manufacturer’s instructions. Cells were analyzed 48 h after transfection.

### Immunostaining and imaging of HeLa cells

Cells grown on coverslips were fixed with 3.7% formaldehyde for 15 min at RT and permeabilized with 0.2% Triton X-100 for 2 min. Cells were blocked with 1% bovine serum albumin for 10 min and incubated with primary antibodies for 1 h at 37°C. After washing with PBS, cells were incubated with secondary antibodies for 1 h at 37°C. After washing with PBS, cells were mounted using VECTASHIELD mounting medium (#H-1000; Vector Laboratories, Inc., CA).

The following primary antibodies were used: mouse anti-acetylated tubulin (1:500; Merck, Darmstadt, Germany) and rat anti-HA (1:500; Roche Diagnostics GmbH). The following secondary antibodies were used: goat anti-mouse IgG Alexa Fluor 633 (1:200; Thermo Fisher Scientific) and goat anti-rat IgG Alexa Fluor 568 (1:250; Thermo Fisher Scientific). ImageJ software (NIH, v.1.54f) was used to quantify the fluorescence intensity.

### Statistical analysis

All data were analyzed and plotted using GraphPad Prism 10 (GraphPad Software, San Diego, CA, USA). Statistical significance was determined using the statistical tests described in the corresponding figure legends. P values of < 0.05 were considered statistically significant.

## REFERENCES

1. Barisic, N., Claeys, K.G., Sirotković-Skerlev, M., Löfgren, A., Nelis, E., De Jonghe, P., and Timmerman, V. (2008). Charcot-Marie-Tooth disease: a clinico-genetic confrontation: Charcot-Marie-tooth disease. Ann. Hum. Genet. 72, 416–441. 10.1111/j.1469-1809.2007.00412.x.

2. Patzkó, A., and Shy, M.E. (2011). Update on Charcot-Marie-Tooth disease. Curr. Neurol. Neurosci. Rep. 11, 78–88. 10.1007/s11910-010-0158-7.

3. Azevedo, H., Pupe, C., Pereira, R., and Nascimento, O.J.M. (2018). Pain in Charcot-Marie-Tooth disease: an update. Arq. Neuropsiquiatr. 76, 273–276. 10.1590/0004-282x20180021.

4. Hattori, N., Yamamoto, M., Yoshihara, T., Koike, H., Nakagawa, M., Yoshikawa, H., Ohnishi, A., Hayasaka, K., Onodera, O., Baba, M., et al. (2003). Demyelinating and axonal features of Charcot-Marie-Tooth disease with mutations of myelin-related proteins (PMP22, MPZ and Cx32): a clinicopathological study of 205 Japanese patients. Brain 126, 134–151. 10.1093/brain/awg012.

5. Suter, U., and Scherer, S.S. (2003). Disease mechanisms in inherited neuropathies. Nat. Rev. Neurosci. 4, 714–726. 10.1038/nrn1196.

6. Züchner, S., Noureddine, M., Kennerson, M., Verhoeven, K., Claeys, K., De Jonghe, P., Merory, J., Oliveira, S.A., Speer, M.C., Stenger, J.E., et al. (2005). Mutations in the pleckstrin homology domain of dynamin 2 cause dominant intermediate Charcot-Marie-Tooth disease. Nat. Genet. 37, 289–294. 10.1038/ng1514.

7. Bitoun, M., Stojkovic, T., Prudhon, B., Maurage, C.-A., Latour, P., Vermersch, P., and Guicheney, P. (2008). A novel mutation in the dynamin 2 gene in a Charcot-Marie-Tooth type 2 patient: clinical and pathological findings. Neuromuscul. Disord. 18, 334–338. 10.1016/j.nmd.2008.01.005.

8. Shpetner, H.S., and Vallee, R.B. (1989). Identification of dynamin, a novel mechanochemical enzyme that mediates interactions between microtubules. Cell 59, 421–432. 10.1007/978-3-540-48711-1_22.

9. Scaife, R., and Margolis, R.L. (1990). Biochemical and immunochemical analysis of rat brain dynamin interaction with microtubules and organelles in vivo and in vitro. J. Cell Biol. 111, 3023–3033. 10.1083/jcb.111.6.3023.

10. Maeda, K., Nakata, T., Noda, Y., Sato-Yoshitake, R., and Hirokawa, N. (1992). Interaction of dynamin with microtubules: its structure and GTPase activity investigated by using highly purified dynamin. Mol. Biol. Cell 3, 1181–1194. 10.1091/mbc.3.10.1181.

11. Shpetner, H.S., and Vallee, R.B. (1992). Dynamin is a GTPase stimulated to high levels of activity by microtubules. Nature 355, 733–735. 10.1038/355733a0.

12. Chen, M.S., Obar, R.A., Schroeder, C.C., Austin, T.W., Poodry, C.A., Wadsworth, S.C., and Vallee, R.B. (1991). Multiple forms of dynamin are encoded by shibire, a Drosophila gene involved in endocytosis. Nature 351, 583–586. 10.1038/351583a0.

13. Clark, S.G., Shurland, D.L., Meyerowitz, E.M., Bargmann, C.I., and van der Bliek, A.M. (1997). A dynamin GTPase mutation causes a rapid and reversible temperature-inducible locomotion defect in C. elegans. Proc. Natl. Acad. Sci. U. S. A. 94, 10438– 10443. 10.1073/pnas.94.19.10438.

14. Cao, H., Garcia, F., and McNiven, M.A. (1998). Differential distribution of dynamin isoforms in mammalian cells. Mol. Biol. Cell 9, 2595–2609. 10.1091/mbc.9.9.2595.

15. Nakata, T., Takemura, R., and Hirokawa, N. (1993). A novel member of the dynamin family of GTP-binding proteins is expressed specifically in the testis. J. Cell Sci. 105 ( *Pt 1*), 1–5. 10.1242/jcs.105.1.1.

16. Sontag, J.M., Fykse, E.M., Ushkaryov, Y., Liu, J.P., Robinson, P.J., and Südhof, T.C. (1994). Differential expression and regulation of multiple dynamins. J. Biol. Chem. 269, 4547–4554. 10.1016/s0021-9258(17)41812-6.

17. Damke, H., Baba, T., Warnock, D.E., and Schmid, S.L. (1994). Induction of mutant dynamin specifically blocks endocytic coated vesicle formation. J. Cell Biol. 127, 915–934. 10.1083/jcb.127.4.915.

18. Warnock, D.E., Baba, T., and Schmid, S.L. (1997). Ubiquitously expressed dynamin-II has a higher intrinsic GTPase activity and a greater propensity for self-assembly than neuronal dynamin-I. Mol. Biol. Cell 8, 2553–2562. 10.1091/mbc.8.12.2553.

19. Tanabe, K., and Takei, K. (2009). Dynamic instability of microtubules requires dynamin 2 and is impaired in a Charcot-Marie-Tooth mutant. J. Cell Biol. 185, 939– 948. 10.1083/jcb.200803153.

20. Guo, R., Fujito, R., Nagasaki, A., Okumura, M., Chihara, T., and Hamao, K. (2022). Dynamin - 2 regulates microtubule stability via an endocytosis - independent mechanism. Cytoskeleton (Hoboken) 79, 94‒104. 10.1002/cm.21723.

21. Levi, T., Sloutskin, A., Kalifa, R., Juven-Gershon, T., and Gerlitz, O. (2020). Efficient in vivo introduction of point mutations using ssODN and a co-CRISPR approach. Biol. Proced. Online 22, 14. 10.1186/s12575-020-00123-7.

22. Pareyson, D., and Marchesi, C. (2009). Diagnosis, natural history, and management of Charcot-Marie-Tooth disease. Lancet Neurol. 8, 654–667. 10.1016/S1474-4422(09)70110-3.

23. Kitani-Morii, F., and Noto, Y.I. (2020). Recent advances in drosophila models of charcot-marie-tooth disease. Int. J. Mol. Sci. 21, 1–19. 10.3390/ijms21197419.

24. Sleigh, J.N., Grice, S.J., Burgess, R.W., Talbot, K., and Cader, M.Z. (2014). Neuromuscular junction maturation defects precede impaired lower motor neuron connectivity in Charcot-Marie-Tooth type 2D mice. Hum. Mol. Genet. 23, 2639– 2650. 10.1093/hmg/ddt659.

25. Morant, L., Erfurth, M.-L., and Jordanova, A. (2021). Drosophila Models for Charcot-Marie-Tooth Neuropathy Related to Aminoacyl-tRNA Synthetases. Genes (Basel) 12, 1519. 10.3390/genes12101519.

26. Hur, J.H., Lee, S.-H., Kim, A.-Y., and Koh, Y.H. (2018). Regulation of synaptic architecture and synaptic vesicle pools by Nervous wreck at Drosophila Type 1b glutamatergic synapses. Exp. Mol. Med. 50, e462. 10.1038/emm.2017.303.

27. Miller, D.L., Ballard, S.L., and Ganetzky, B. (2012). Analysis of synaptic growth and function in Drosophila with an extended larval stage. J. Neurosci. 32, 13776–13786. 10.1523/JNEUROSCI.0508-12.2012.

28. O’Connor-Giles, K.M., and Ganetzky, B. (2008). Satellite signaling at synapses. Fly (Austin) 2, 259–261. 10.4161/fly.7133.

29. Chou, V.T., Johnson, S.A., and Van Vactor, D. (2020). Synapse development and maturation at the drosophila neuromuscular junction. Neural Dev. 15, 11. 10.1186/s13064-020-00147-5.

30. Bodaleo, F.J., and Gonzalez-Billault, C. (2016). The presynaptic microtubule cytoskeleton in physiological and pathological conditions: Lessons from Drosophila fragile X syndrome and hereditary spastic paraplegias. Front. Mol. Neurosci. 9, 60. 10.3389/fnmol.2016.00060.

31. Coombes, C.E., Saunders, H.A.J., Mannava, A.G., Johnson-Schlitz, D.M., Reid, T.A., Parmar, S., McClellan, M., Yan, C., Rogers, S.L., Parrish, J.Z., et al. (2020). Non-enzymatic activity of the α-tubulin acetyltransferase αTAT limits synaptic Bouton growth in neurons. Curr. Biol. 30, 610–623.e5. 10.1016/j.cub.2019.12.022.

32. Saunders, H.A.J., Johnson-Schlitz, D.M., Jenkins, B.V., Volkert, P.J., Yang, S.Z., and Wildonger, J. (2022). Acetylated α-tubulin K394 regulates microtubule stability to shape the growth of axon terminals. Curr. Biol. 32, 614–630.e5. 10.1016/j.cub.2021.12.012.

33. Palazzo, A., Ackerman, B., and Gundersen, G.G. (2003). Cell biology: Tubulin acetylation and cell motility: Cell biology (Communication arising). Nature 421, 230. 10.1038/421230a.

34. Hummel, T., Krukkert, K., Roos, J., Davis, G., and Klämbt, C. (2000). Drosophila Futsch/22C10 is a MAP1B-like protein required for dendritic and axonal development. Neuron 26, 357–370. 10.1016/s0896-6273(00)81169-1.

35. Prokop, A. (2020). Cytoskeletal organization of axons in vertebrates and invertebrates. J. Cell Biol. 219. 10.1083/jcb.201912081.

36. Roos, J., Hummel, T., Ng, N., Klämbt, C., and Davis, G.W. (2000). Drosophila Futsch regulates synaptic microtubule organization and is necessary for synaptic growth. Neuron 26, 371–382. 10.1016/s0896-6273(00)81170-8.

37. Packard, M., Koo, E.S., Gorczyca, M., Sharpe, J., Cumberledge, S., and Budnik, V. (2002). The Drosophila Wnt, wingless, provides an essential signal for pre- and postsynaptic differentiation. Cell 111, 319–330. 10.1016/s0092-8674(02)01047-4.

38. Sherwood, N.T., Sun, Q., Xue, M., Zhang, B., and Zinn, K. (2004). Drosophila spastin regulates synaptic microtubule networks and is required for normal motor function. PLoS Biol. 2, e429. 10.1371/journal.pbio.0020429.

39. Vasquez, R.J., Howell, B., Yvon, A.M., Wadsworth, P., and Cassimeris, L. (1997). Nanomolar concentrations of nocodazole alter microtubule dynamic instability in vivo and in vitro. Mol. Biol. Cell 8, 973–985. 10.1091/mbc.8.6.973.

40. Yvon, A.M., Wadsworth, P., and Jordan, M.A. (1999). Taxol suppresses dynamics of individual microtubules in living human tumor cells. Mol. Biol. Cell 10, 947–959. 10.1091/mbc.10.4.947.

41. Xiao, H., Verdier-Pinard, P., Fernandez-Fuentes, N., Burd, B., Angeletti, R., Fiser, A., Horwitz, S.B., and Orr, G.A. (2006). Insights into the mechanism of microtubule stabilization by Taxol. Proc. Natl. Acad. Sci. U. S. A. 103, 10166–10173. 10.1073/pnas.0603704103.

42. Baines, A.J., Bignone, P.A., King, M.D.A., Maggs, A.M., Bennett, P.M., Pinder, J.C., and Phillips, G.W. (2009). The CKK domain (DUF1781) binds microtubules and defines the CAMSAP/ssp4 family of animal proteins. Mol. Biol. Evol. 26, 2005– 2014. 10.1093/molbev/msp115.

43. Goodwin, S.S., and Vale, R.D. (2010). Patronin Regulates the Microtubule Network by Protecting Microtubule Minus Ends. Cell 143, 263–274. 10.1016/j.cell.2010.09.022.

44. Wang, H., Brust-Mascher, I., Civelekoglu-Scholey, G., and Scholey, J.M. (2013). Patronin mediates a switch from kinesin-13-dependent poleward flux to anaphase B spindle elongation. J. Cell Biol. 203, 35–46. 10.1083/jcb.201306001.

45. Hendershott, M.C., and Vale, R.D. (2014). Regulation of microtubule minus-end dynamics by CAMSAPs and Patronin. Proc. Natl. Acad. Sci. U. S. A. 111, 5860– 5865. 10.1073/pnas.1404133111.

46. Martin, M., and Akhmanova, A. (2018). Coming into focus: Mechanisms of microtubule minus-end organization. Trends Cell Biol. 28, 574–588. 10.1016/j.tcb.2018.02.011.

47. Yang, D.W., and Choi, K.W. (2021). Suppression of Patronin deficiency by altered Hippo signaling in Drosophila organ development. Cell Death Differ. 28, 233–250. 10.1038/s41418-020-0597-x.

48. Sidiropoulos, P.N.M., Miehe, M., Bock, T., Tinelli, E., Oertli, C.I., Kuner, R., Meijer, D., Wollscheid, B., Niemann, A., and Suter, U. (2012). Dynamin 2 mutations in Charcot-Marie-Tooth neuropathy highlight the importance of clathrin-mediated endocytosis in myelination. Brain 135, 1395–1411. 10.1093/brain/aws061.

49. Menon, K.P., Carrillo, R.A., and Zinn, K. (2013). Development and plasticity of the Drosophila larval neuromuscular junction: Development and plasticity of the neuromuscular junction. Wiley Interdiscip. Rev. Dev. Biol. 2, 647–670. 10.1002/wdev.108.

50. Dickman, D.K., Lu, Z., Meinertzhagen, I.A., and Schwarz, T.L. (2006). Altered synaptic development and active zone spacing in endocytosis mutants. Curr. Biol. 16, 591–598. 10.1016/j.cub.2006.02.058.

51. Jiang, K., Faltova, L., Hua, S., Capitani, G., Prota, A.E., Landgraf, C., Volkmer, R., Kammerer, R.A., Steinmetz, M.O., and Akhmanova, A. (2018). Structural Basis of Formation of the Microtubule Minus-End-Regulating CAMSAP-Katanin Complex. Structure 26, 375–382.e4. 10.1016/j.str.2017.12.017.

52. Ferguson, S.M., and De Camilli, P. (2012). Dynamin, a membrane-remodelling GTPase. Nat. Rev. Mol. Cell Biol. 13, 75–88. 10.1038/nrm3266.

53. Tassin, T.C., Barylko, B., Hedde, P.N., Chen, Y., Binns, D.D., James, N.G., Mueller, J.D., Jameson, D.M., Taussig, R., and Albanesi, J.P. (2021). Gain-of-Function Properties of a Dynamin 2 Mutant Implicated in Charcot-Marie-Tooth Disease. Front. Cell. Neurosci. 15, 1–10. 10.3389/fncel.2021.745940.

54. Castells-Nobau, A., Nijhof, B., Eidhof, I., Wolf, L., Scheffer-de Gooyert, J.M., Monedero, I., Torroja, L., van der Laak, J.A.W.M., and Schenck, A. (2017). Two algorithms for high-throughput and multi-parametric quantification of Drosophila neuromuscular junction morphology. J. Vis. Exp. 10.3791/55395.

