## Supplemental Information for "A Charcot–Marie–Tooth–associated dynamin mutation causes peripheral neuropathy-like phenotypes via Patronin/CAMSAP-dependent microtubule stabilization"

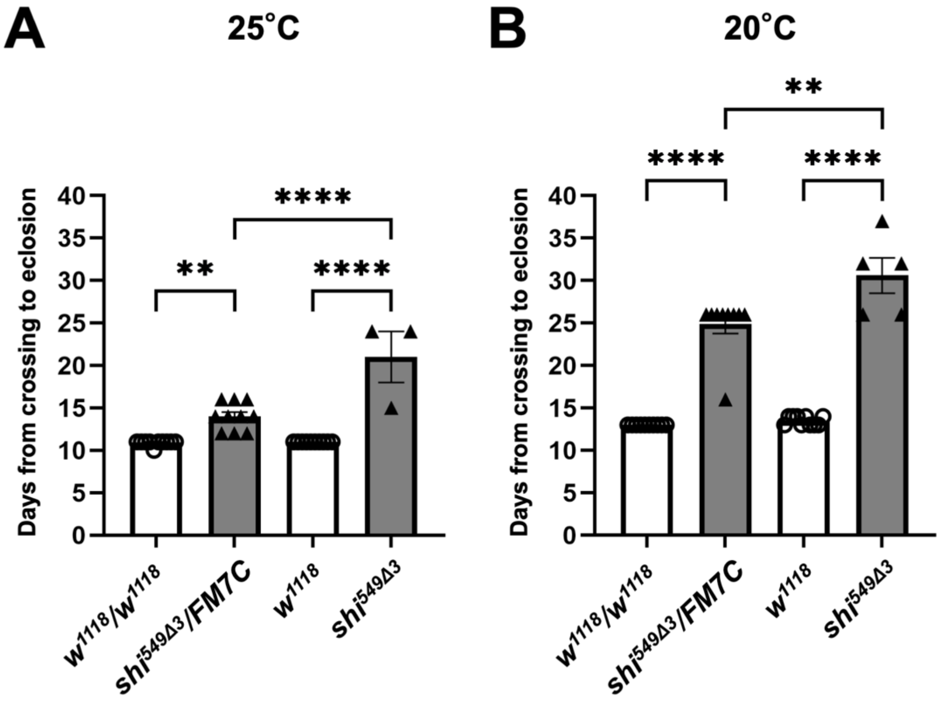


**Figure S1. Prolonged developmental time in *shi*$^{549\Delta3}$ flies**

(A, B) Time from egg laying (crossing) to eclosion at 25°C (A) and 20°C (B) in control and *shi*$^{549\Delta3}$ flies. Control flies were generated by crossing *w*$^{1118}$/*w*$^{1118}$ females with *w*$^{1118}$ males, whereas *shi*$^{549\Delta3}$ mutant flies were generated by crossing *shi*$^{549\Delta3}$/FM7C females with FM7C males. The following genotypes were analyzed: control females (*w*$^{1118}$/*w*$^{1118}$), *shi*$^{549\Delta3}$ females (*shi*$^{549\Delta3}$/FM7C), control males (*w*$^{1118}$), and *shi*$^{549\Delta3}$ males (*shi*$^{549\Delta3}$). At 25°C, n = 10 (*w*$^{1118}$/*w*$^{1118}$), n = 10 (*shi*$^{549\Delta3}$/FM7C), n = 10 (*w*$^{1118}$), and n = 3 (*shi*$^{549\Delta3}$). At 20°C, n = 10 (*w*$^{1118}$/*w*$^{1118}$), n = 9 (*shi*$^{549\Delta3}$/FM7C), n = 10 (*w*$^{1118}$), and n = 5 (*shi*$^{549\Delta3}$). Statistical analysis was performed using unpaired t-tests (**P < 0.01, ****P < 0.0001). Error bars indicate mean ± s.e.m.

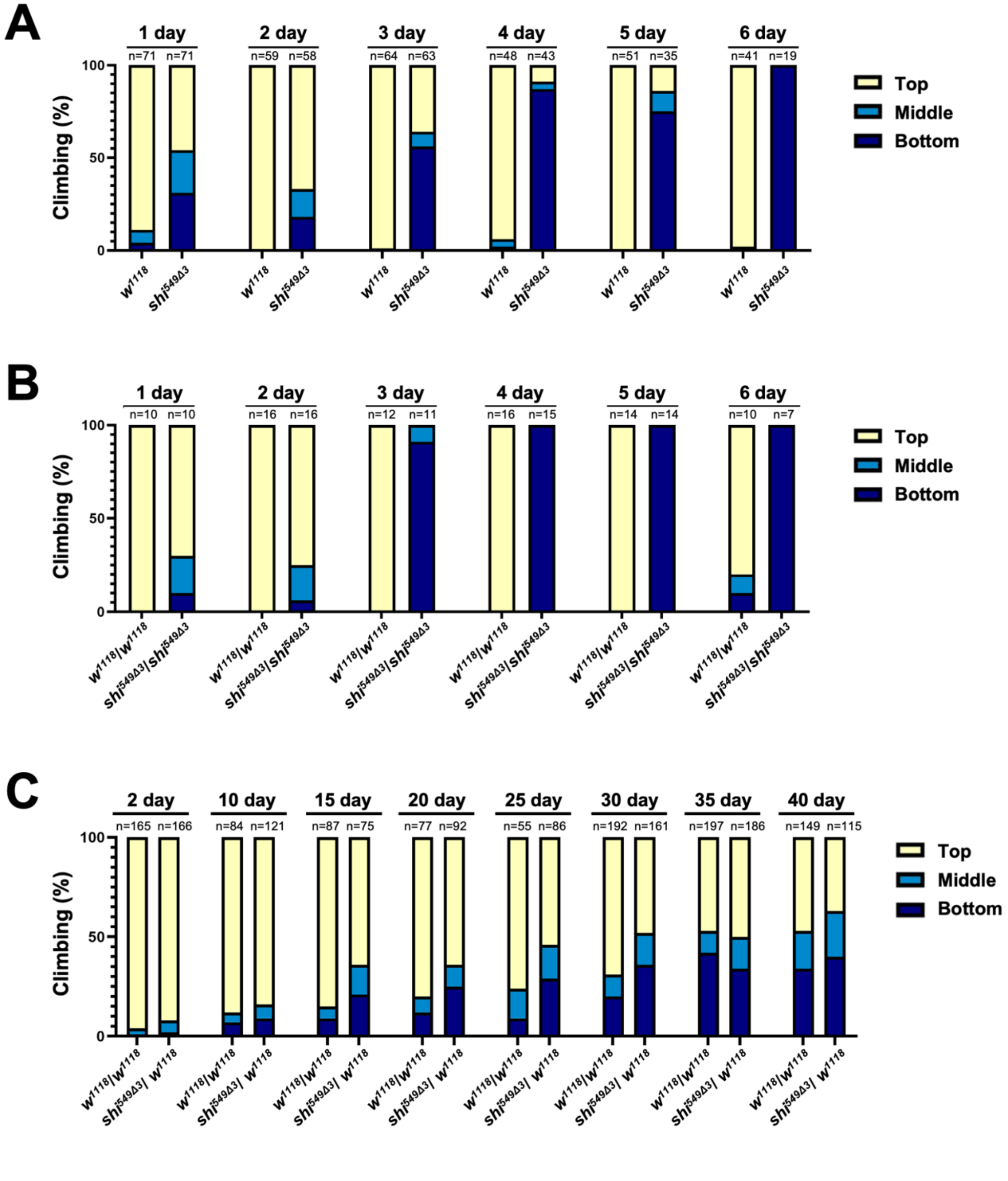


**Figure S2.** ***shi*$^{549\Delta3}$ mutant flies exhibit progressive motor impairment**

(A–C) Forced climbing assays showing the detailed distribution of flies reaching the top (yellow), middle (light blue), or remaining at the bottom (dark blue) of the vial for control ($w^{1118}$) and $shi^{549\Delta3}$ mutant adult flies: (A) hemizygous ($shi^{549\Delta3}$), (B) homozygous ($shi^{549\Delta3}/shi^{549\Delta3}$), and (C) heterozygous ($shi^{549\Delta3}/w^{1118}$). Numbers above each graph indicate age (days post-eclosion). Sample sizes are indicated in the graphs.

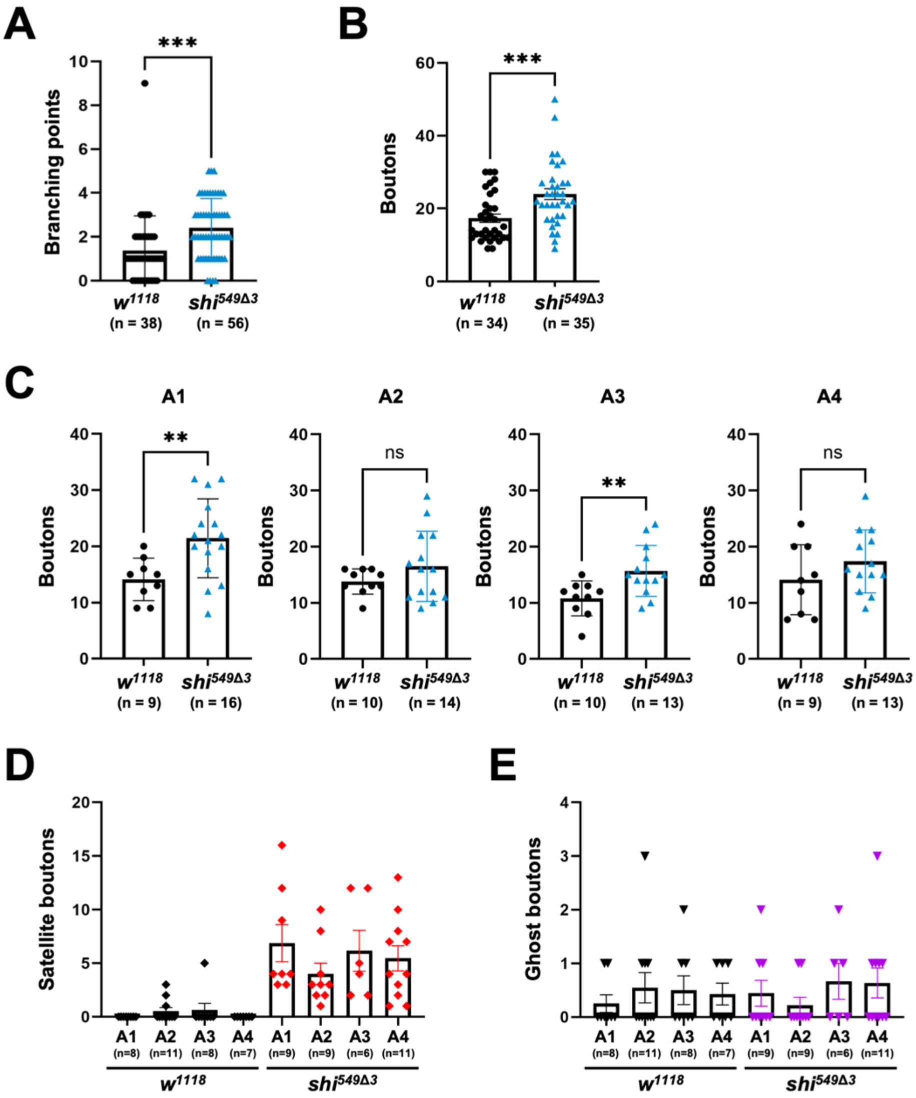


**Figure S3. Altered neuromuscular junction (NMJ) morphology in *shi*$^{549\Delta3}$ mutant larvae**

(A) Quantification of axonal branching points (initiation sites of axonal branches) at NMJs on muscle 4 in abdominal segments A1–A4 from five control (*w*$^{1118}$) larvae (n = 38 NMJs) and 11 *shi*$^{549\Delta3}$ mutant larvae (n = 56 NMJs).

(B) Quantification of type Ib boutons (excluding ectopic boutons) at NMJs on muscle 4 in abdominal segments A1–A4 from five control (*w*$^{1118}$) larvae (n = 34 NMJs) and eight *shi*$^{549\Delta3}$ mutant larvae (n = 35 NMJs).

(C–E) Quantification of type Ib (C), satellite (D), and ghost boutons (E) per muscle segment (A1–A4). NMJs from five control larvae and 11 *shi*$^{549\Delta3}$ mutant larvae were analyzed for (C), and NMJs from five control larvae and eight *shi*$^{549\Delta3}$ mutant larvae were analyzed for (D–E). The number of NMJs analyzed for each dataset is indicated in the graphs.

Statistical analysis was performed using multiple unpaired t-tests (*P < 0.05, **P < 0.01, ***P < 0.001, ****P < 0.0001; ns, not significant). Error bars indicate mean ± s.e.m.

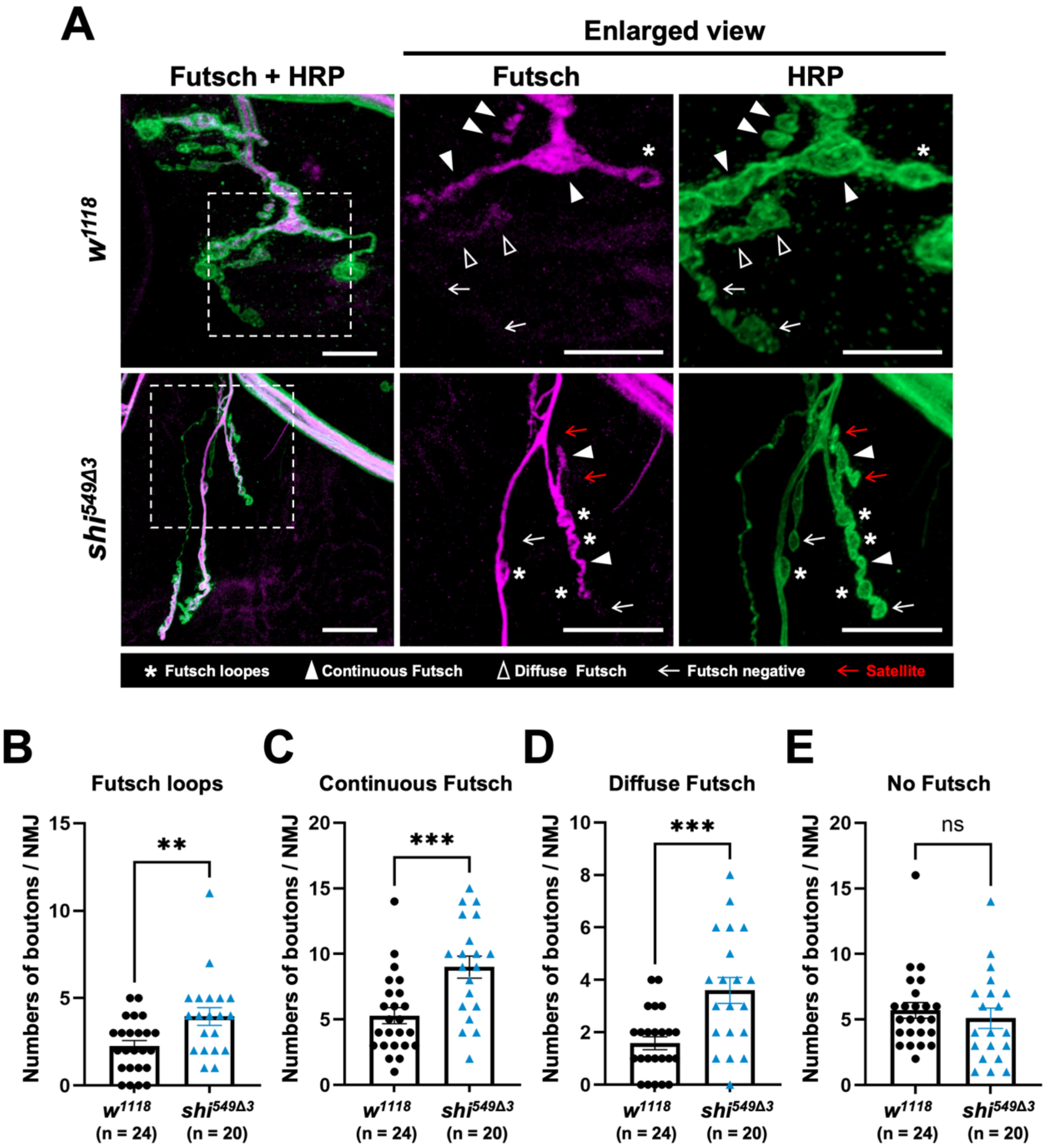


**Figure S4.** ***shi***$^{549\Delta3}$ **mutant flies show altered Futsch staining patterns**

(A) Representative images of larval NMJs from control (*w*$^{1118}$) and hemizygous *shi*$^{549\Delta3}$ mutant larvae stained with antibodies against Futsch (magenta) and horseradish peroxidase (HRP; green). Futsch staining patterns at NMJs on muscle 4 (A1–A4) were classified as loop (asterisks), continuous (bundles or splayed bundles; filled triangles), diffuse (open triangles), or negative (undetectable; arrows) for each bouton. Scale bar, 20 μm. (B–E) Quantification of the number of boutons per NMJ on muscle 4 (A1–A4), excluding satellite boutons, according to the Futsch staining patterns described above. NMJs from five control larvae (n = 24) and five *shi*$^{549\Delta3}$ mutant larvae (n = 20) were analyzed. Statistical analysis was performed using unpaired t-tests (**P < 0.01, ***P < 0.001; ns, not significant). Error bars indicate mean ± s.e.m.
